# Post activation potentiation is greater in human triceps brachii versus triceps surae muscles

**DOI:** 10.1101/2025.04.23.650262

**Authors:** William S. Zoughaib, Madison J. Fry, Ahaan Singhal, Richard L. Hoffman, Andrew R. Coggan

**Author notes:** Address correspondence to: Andrew R. Coggan, PhD, FACSM, Department of Kinesiology, Indiana University Indianapolis, IF 101C, 250 University Boulevard, Indianapolis, IN 46112.

## Abstract

**Introduction/Aims:** Post activation potentiation (PAP) is an inherent characteristic of muscle wherein an increase in twitch force is observed after voluntary activation. This phenomenon is believed to be largely due to phosphorylation of the regulatory light chain of myosin. In rodents and other species, PAP occurs predominantly or exclusively in fast twitch muscles. However, it has been suggested that in humans PAP occurs more or less independently of muscle fiber type.

**Methods:** 18 healthy men and women (27±8 y) underwent an electrical muscle stimulation protocol during which two sets of four twitches were elicited both pre and post 6 s maximal voluntary contractions of both the triceps surae (60-70% slow twitch) and triceps brachii (60-70% fast twitch) muscles.

**Results:** Unpotentiated peak twitch torque (PTT) was higher in the larger triceps surae vs. the smaller triceps brachii (i.e., 13.4±5.3 vs. 3.4±2.1 Nm; P = 8.22 × 10^−13^), but time to peak torque was shorter (i.e., 84±7 vs. 132±14 ms; P = 6.92 × 10^−14^) and the relative rate of torque development (RTD) was greater in the triceps brachii (2294±257 vs. 1425±102 %/s; P = 1.93 × 10^−11^). PAP increased PTT and the absolute RTD by 172±124% and 240±170% in triceps brachii versus only 20±20% and 31±24% in triceps surae, respectively (P = 1.26 × 10^−4^ and 1.30 × 10^−4^, respectively). However, PAP-induced changes in half-relaxation time and the relative rate of relaxation did not differ between the two muscle groups.

**Discussion:** We conclude that PAP influences contraction but not relaxation of human muscle in a fiber type dependent manner.

## Introduction

Post-activation potentiation (PAP) is the short-term (i.e., seconds to minutes) enhancement of skeletal muscle function following prior voluntary contractile activity. Analogous to the post-tetanic potentiation observed following electrically evoked contractions, PAP results in increases in the peak force and rate of force development during an isometric twitch (1). Even greater increases in force (and hence power) are observed during concentric contractions (1). It has been hypothesized that PAP may have evolved as a way of maximizing the neuromuscular efficiency (i.e., work output relative to stimulation rate) (2) and/or economy (i.e., work output relative to energetic cost) (3) of muscle during sustained activity, during which fatigue may otherwise occur. As emphasized by Laidlaw and Vandenboom (2), the physiological significance of PAP is illustrated by the fact that it is observed in numerous species ranging from insects (4) to humans (5).

The increases in force and rate of force development characteristic of PAP are thought to be largely, albeit not entirely, the result of a Ca^2+^-stimulated increase in phosphorylation of the myosin regulatory light chain (RLC) (6). This disrupts the superrelaxed state of myosin, in which the myosin heads interact with each other and are folded back against the myosin filament (7). The resulting increase in the proportion of “swaying” myosin heads that are free to bind with actin leads to an increase in Ca^2+^ sensitivity and thus force upon stimulation. Furthermore, although RLC phosphorylation has no apparent effect on the maximal ATPase activity or velocity of unloaded shortening of muscle (1), which are indicative of the rate of crossbridge cycling, the formation of more crossbridges results in an increase in the rate of force development when Ca^2+^ levels are subsaturating (e.g., during a twitch) (8).

In fish (9), birds (10), cats (11), rats (12), rabbits (13), and mice (14), significant PAP is observed only in fast and not in slow muscles. At least in rodents, this appears to be primarily due to a >10-fold higher ratio of skeletal muscle myosin light chain kinase (smMLCK) activity to myosin light chain phosphatase activity in fast muscle fibers (15). As a result, during contractions a marked increase in RLC phosphorylation occurs only in fast fibers, and in fact it has been suggested that the potentiated state is the normal operating state of fast muscles (16). In humans, however, muscle contraction results in phosphorylation of the RLC in both fiber types (17,18), and considerable PAP is observed even in muscles with a high percentage of slow fibers, e.g., the plantar flexors (19-24). Moreover, endurance athletes exhibit greater PAP in their trained muscles despite a slower phenotype (24). Contrastingly, PAP has also been reported to be greater in faster vs. slower human muscles (23,25) and to be inversely correlated with baseline (i.e., unpotentiated) twitch kinetics (25,26), implying a fast twitch fiber dependence. Thus, the effects of muscle fiber type on PAP in humans are still uncertain (1).

In addition to altering the rate of force development, PAP may (or may not) alter the rate of relaxation. In rodent muscle, PAP slows the rate of relaxation from a steady force but enhances the rate of relaxation during a twitch (1), effects that appear to be independent of either RLC phosphorylation or muscle fiber type (26). In humans, on the other hand, PAP has been reported to either reduce (20,22,24,28), not change (20,21,26,29), or even increase (30) twitch half-relaxation time in various muscle groups. The effects of PAP on relaxation of human muscle, especially with respect to fiber type, are therefore still unclear.

The purpose of the present study was to determine the effects of PAP on the twitch contractile properties of two human muscles differing significantly in fiber type composition, i.e., the triceps brachii (60-70% fast twitch) and triceps surae (60-70% slow twitch) muscles (31,32). We hypothesized that PAP would increase twitch force and kinetics in both muscle groups, but that these effects would be larger in the triceps brachii vs. the triceps surae.

## Methods

### Participants

After providing written informed consent, 18 normally active men and women completed this study, which was approved by the Human Subjects Office at Indiana University. The characteristics of these participants are shown in Table 1. To avoid the potentially confounding influence of exercise training on PAP (24), highly active individuals (i.e., those performing >3000 MET-min/wk of moderate physical activity or >1500 MET-min/wk of vigorous physical activity based on the short-form International Physical Activity Questionnaire (33)) were excluded. Other exclusion criteria were age <18 or >44 y, current use of antibiotics, phosphodiesterase inhibitors, tobacco, or any supplements intended to increase muscle mass or function, diagnosis of epilepsy or presence of a pacemaker or other implantable cardiac device, resting blood pressure >140/90 mmHg, or an answer of yes to any of the seven general health questions of the Physical Activity Readiness Questionnaire (34). In addition, women were excluded if they had an irregular menstrual cycle (average length <21 or >35 d), had missed more than three consecutive periods in the last 12 mo, or were pregnant, using hormonal contraceptives, or on hormone replacement therapy. We did not attempt to control for menstrual cycle phase since it has been reported to have no influence on twitch contractile properties (35,36).

**Table 1.** Participant characteristics.

|  |  |
| --- | --- |
| Sex (M/F) | 15/3 |
| Age (y) | 27 ± 8 |
| Height (m) | 1.74 ± 0.07 |
| Mass (kg) | 77.0 ± 18.3 |
| BMI (kg/m<sup>2</sup>) | 25.2 ± 4.8 |
| IPAQ Score (MET-min/wk) | 1140 ± 751 |

BMI, body mass index. IPAQ, International Physical Activity Questionnaire. Values are mean ± SD for n=18.

### Experimental procedures

A constant current stimulator (Digitimer DS 7AH, Hertfordshire, UK) was used to elicit unpotentiated and potentiated isometric twitch contractions of the triceps brachii and triceps surae, with the resultant torque measured at 1000 Hz using an isokinetic dynamometer (Biodex System 4 Pro, Biodex Medical Systems, Shirley, NY) and data acquisition system (Biopac MP160, Biopac, Goleta, CA). The order of testing of the two muscle groups was randomized between participants.

After shaving, abrading, and cleaning the participant’s skin over the muscles of interest, two 5.08 cm x 8.89 cm self-adhesive electrodes (Dura-Stick Plus, Chattanooga Medical Supply, Chattanooga, TN) were applied to the first muscle to be tested. For the triceps brachii, the cathode was placed diagonally across the proximal posterolateral portion of the long and lateral heads, whereas the anode was placed transversely ~5 cm proximal to the olecranon process. The participant was then positioned on the dynamometer with a chair back angle of 1.48 rad (85°) and with the shoulder of the dominant arm adducted and at an elevation of 0 rad (0°). The elbow was supported at a joint angle of 1.57 rad (90°) and straps were placed over the participant’s waist, torso, and forearm to restrict extraneous movement. A plastic and fabric brace (Roylan Ulnar Deviation Splint, Performance Health, Cedarburg, WI) and an elastic wrap were used to prevent any movement of the wrist or fingers relative to the handgrip. For the triceps surae, the cathode was placed transversely over the gastrocnemius muscle ~5 cm distal to the popliteal fossa, whereas the anode was placed over the soleus muscle ~10 cm proximal to the calcaneus. The participant was then positioned on the dynamometer with a chair back angle of 0.96 rad (55°) and the knee and ankle joints of the ipsilateral leg of the dominant arm at an angle of 0 rad (0°). Straps were placed over the waist and thigh as well as across the instep and metatarsals to prevent any movement of the leg or the foot with respect to the footplate.

Once the participant was positioned, single stimuli (400 V, 200 µs) of increasing current were applied at 3-5 s intervals until a plateau in twitch torque was observed. After a brief rest, four unpotentiated twitches were then elicited at approximately 1 s intervals. The participant then performed a 6 s maximal voluntary isometric contraction (MVC), during which strong verbal encouragement was provided. Immediately afterwards, four potentiated twitches were elicited, also at approximately 1 s intervals. Results of a representative experiment are shown in Fig. 1. After 10 min of rest (19,29), a single stimulus was applied to verify reversal of the potentiation, then this sequence was repeated. The electrodes were then moved, and the participant repositioned on the dynamometer to permit testing of the second muscle using the same procedures.

**Figure 1.**
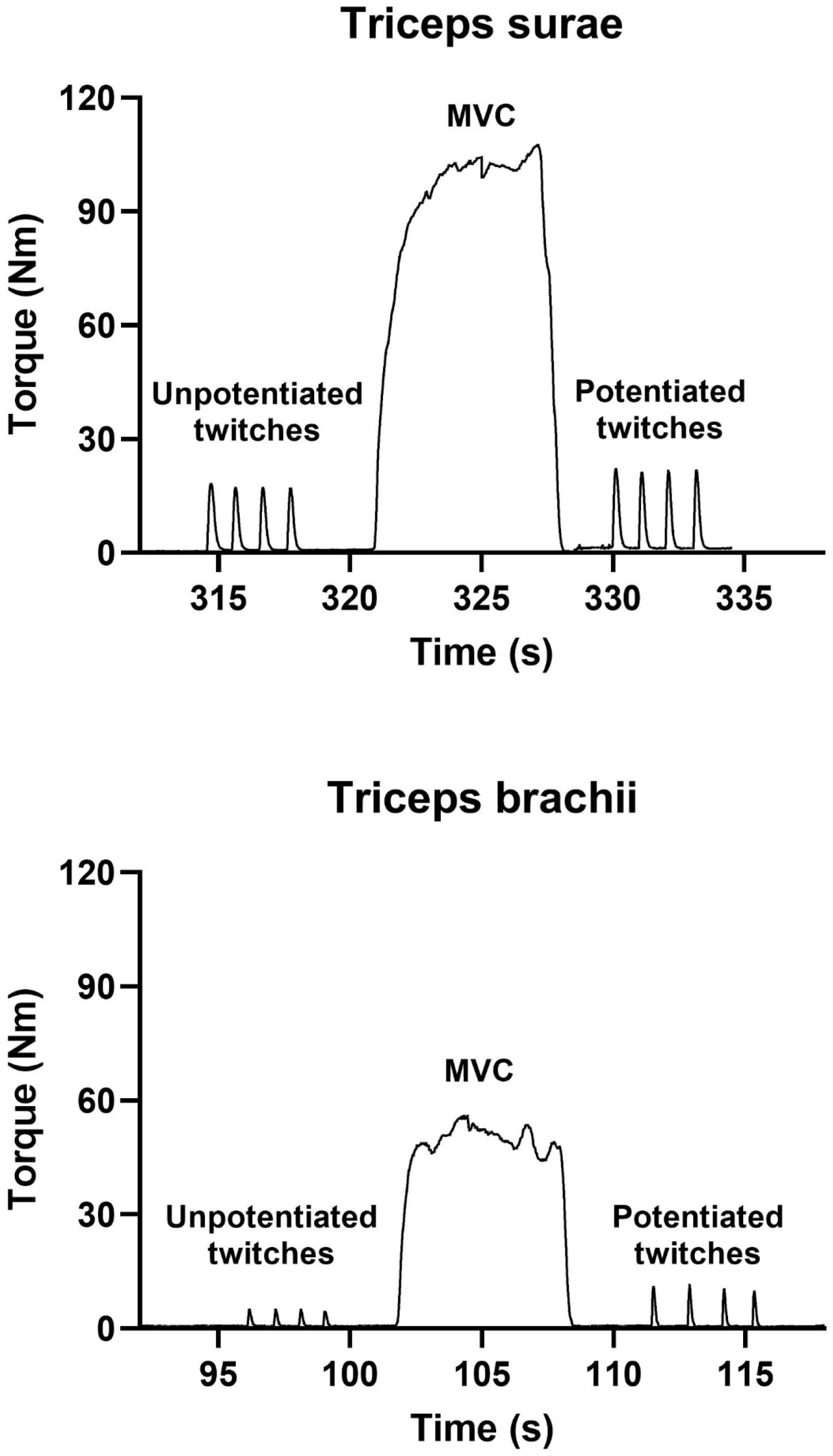
Effects of a 6 s maximal voluntary contraction (MVC) on twitch torque in the triceps surae (*top panel*) or triceps brachii (*bottom panel*) of a representative participant.

### Data analysis

Torque data were analyzed using Biopac AcqKnowledge version 5.08. The signal was first smoothed using a 10 ms rolling average filter, after which peak twitch torque (PTT), time to peak torque (TPT), and one-half relaxation time (HRT) were manually extracted for each twitch. Maximal rates of torque development (RTD) and relaxation (RR) were also determined based on the first derivative of the smoothed torque signal over a 20 ms interval. The eight values obtained for each of these parameters under each condition (i.e., unpotentiated and potentiated) were then averaged and these averages used in all subsequent calculations.

### Statistical analysis

Statistical analyses were performed using GraphPad Prism version 10.4.1 (627) (GraphPad Software, La Jolla, CA). Normality of data distribution was assessed using the D’Agostino and Pearson omnibus test. Two-way (i.e., muscle x condition) analyses of variance (ANOVA) were used to compare twitch contractile properties between the triceps surae and triceps brachii. Differences between individual cell means were tested using the Holm-Šidák multiple comparison procedure. The effects of PAP on percentage changes in PTT, RTD, and RR in the two muscle groups were compared using paired t-tests. Linear regression was used to evaluate the relationship between unpotentiated twitch characteristics and the extent of PAP. Multiplicity-adjusted two-tailed P values <0.05 were considered statistically significant.

## Results

The contractile properties of the two muscle groups are summarized in Table 2. In the unpotentiated state, the PTT of the larger triceps surae was significantly greater than that of the smaller triceps brachii (i.e., P = 8.22 × 10^−13^), whereas TPT and HRT were significantly longer (i.e., P = 6.92 × 10^−14^ and 5.29 × 10^−10^, respectively). Absolute RTD and RR were also greater in the triceps surae (i.e., P = 1.34 × 10^−7^ and 4.19 × 10^−10^, respectively). However, relative RTD and RR were significantly higher in the triceps brachii (i.e., P = 1.93 × 10^−11^ and 9.93 × 10^−6^, respectively).

**Table 2.** Twitch contractile properties of the triceps surae and triceps brachii muscle groups.

|  | Triceps surae |  | Triceps brachii |  | P value |  |  |
| --- | --- | --- | --- | --- | --- | --- | --- |
|  | <u>Unpotentiated</u> | <u>Potentiated</u> | <u>Unpotentiated</u> | <u>Potentiated</u> | <u>Muscle</u> | <u>Condition</u> | <u>Interaction</u> |
| PTT<br>(Nm) | 13.4<br>±5.3 | 15.8<br>±5.9 | 3.4<br>±2.1 | 7.6<br>±2.9 | **2.61 x 10<sup>-7</sup>** | **8.95 x 10<sup>-9</sup>** | **1.85 x 10<sup>-2</sup>** |
| TPT<br>(ms) | 132<br>±14 | 117<br>±9 | 84<br>±7 | 76<br>±9 | **6.27 x 10<sup>-12</sup>** | **8.48 x 10<sup>-7</sup>** | **2.14 x 10<sup>-2</sup>** |
| HRT<br>(ms) | 99<br>±13 | 96<br>±15 | 63<br>±13 | 58<br>±8 | **2.31 x 10<sup>-8</sup>** | **3.72 x 10<sup>-2</sup>** | 0.610 |
| Absolute<br>RTD<br>(Nm/s) | 191<br>±74 | 244<br>±96 | 78<br>±50 | 216<br>±90 | **6.85 x 10<sup>-4</sup>** | **1.10 x 10<sup>-9</sup>** | **1.07 x 10<sup>-4</sup>** |
| Absolute<br>RR<br>(Nm/s) | -111<br>±45 | -131<br>±50 | -41<br>±28 | -94<br>±40 | **1.21 x 10<sup>-5</sup>** | **5.07 x 10<sup>-7</sup>** | **2.23 x 10<sup>-4</sup>** |
| Relative<br>RTD<br>(%/s) | 1425<br>±102 | 1539<br>±137 | 2294<br>±257 | 2850<br>±350 | **1.96 x 10<sup>-11</sup>** | **5.56 x 10<sup>-9</sup>** | **1.28 x 10<sup>-5</sup>** |
| Relative<br>RR<br>(%/s) | -829<br>±82 | -832<br>±105 | -1271<br>±347 | -1241<br>±196 | **1.67 x 10<sup>-6</sup>** | 0.734 | 0.706 |

PTT, peak twitch torque. TPT, time to peak torque. HRT, half relaxation time. RTD, rate of torque development. RR, rate of relaxation. Values are mean ± SD for N=18. P values <0.05 are shown in **bold**.

Potentiation resulted in a significant increase in PTT in both muscle groups, but this effect was larger in the triceps brachii, resulting in a significant muscle x condition interaction effect by ANOVA (Table 2). Nonetheless, the PTT of the triceps surae remained higher that of the triceps brachii (i.e., P = 1.58 × 10^−11^). Potentiation also shortened TPT in both muscle groups, especially in the triceps surae, but the difference between them remained significant (i.e., P = 7.83 × 10^−13^). HRT was also slightly but significantly shorter in both muscles in the potentiated state, but still longer in the triceps surae versus the triceps brachii (i.e., P = 2.90 × 10^−10^). Absolute RTD and RR were accelerated to a greater degree by potentiation in the triceps brachii versus the triceps surae, but remained larger in the triceps surae (i.e., P = 0.0559 and P = 2.66 × 10^−6^, respectively). The relative RTD also increased more in the triceps brachii, magnifying the significance of the difference between the two muscles (i.e., P = 2.88 × 10^−14^ versus 1.93 × 10^−11^ – see above). On the other hand, PAP had no impact on the relative RR in either muscle, such that it remained greater in the triceps brachii (i.e., P = 1.65 × 10^−5^).

Percentage changes in PTT and absolute RTD and RR due to PAP are illustrated in Fig. 2. As shown, the effects of potentiation on these parameters were, on average, almost 10-fold larger in the triceps brachii versus the triceps surae. However, the magnitudes of these changes were not related to baseline, i.e., unpotentiated, twitch kinetics on an individual basis (Fig. 3).

**Figure 2.**
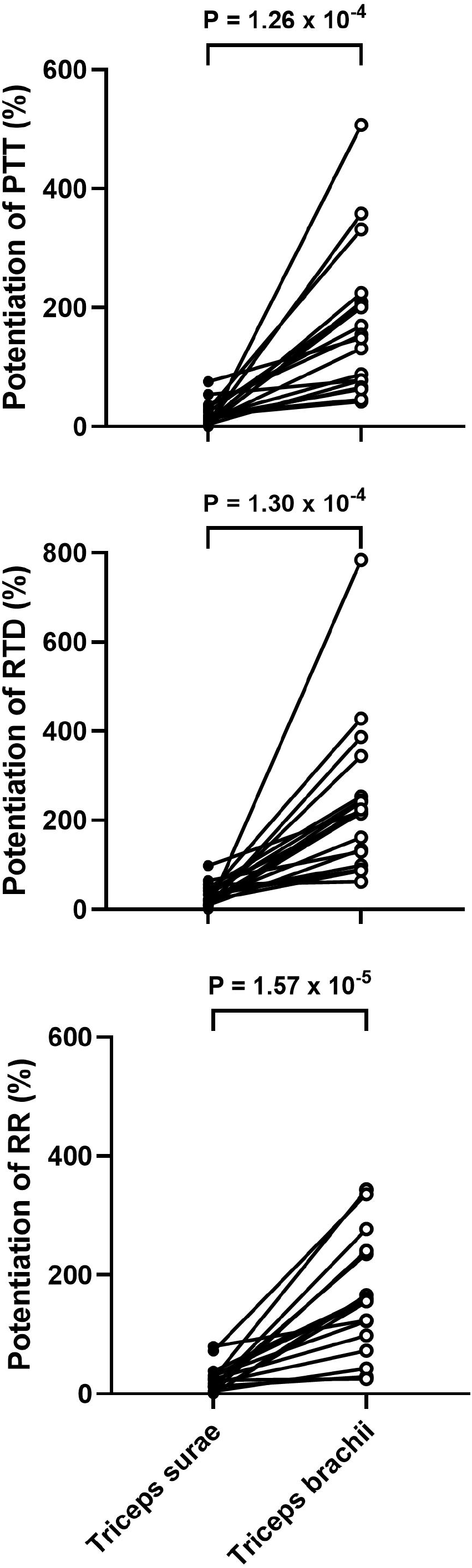
Relative changes in peak twitch torque (PTT, *top panel*), rate of torque development (RTD, *middle* panel), and rate of relaxation (RR, *bottom panel*) due to potentiation in the triceps surae and triceps brachii.

**Figure 3.**
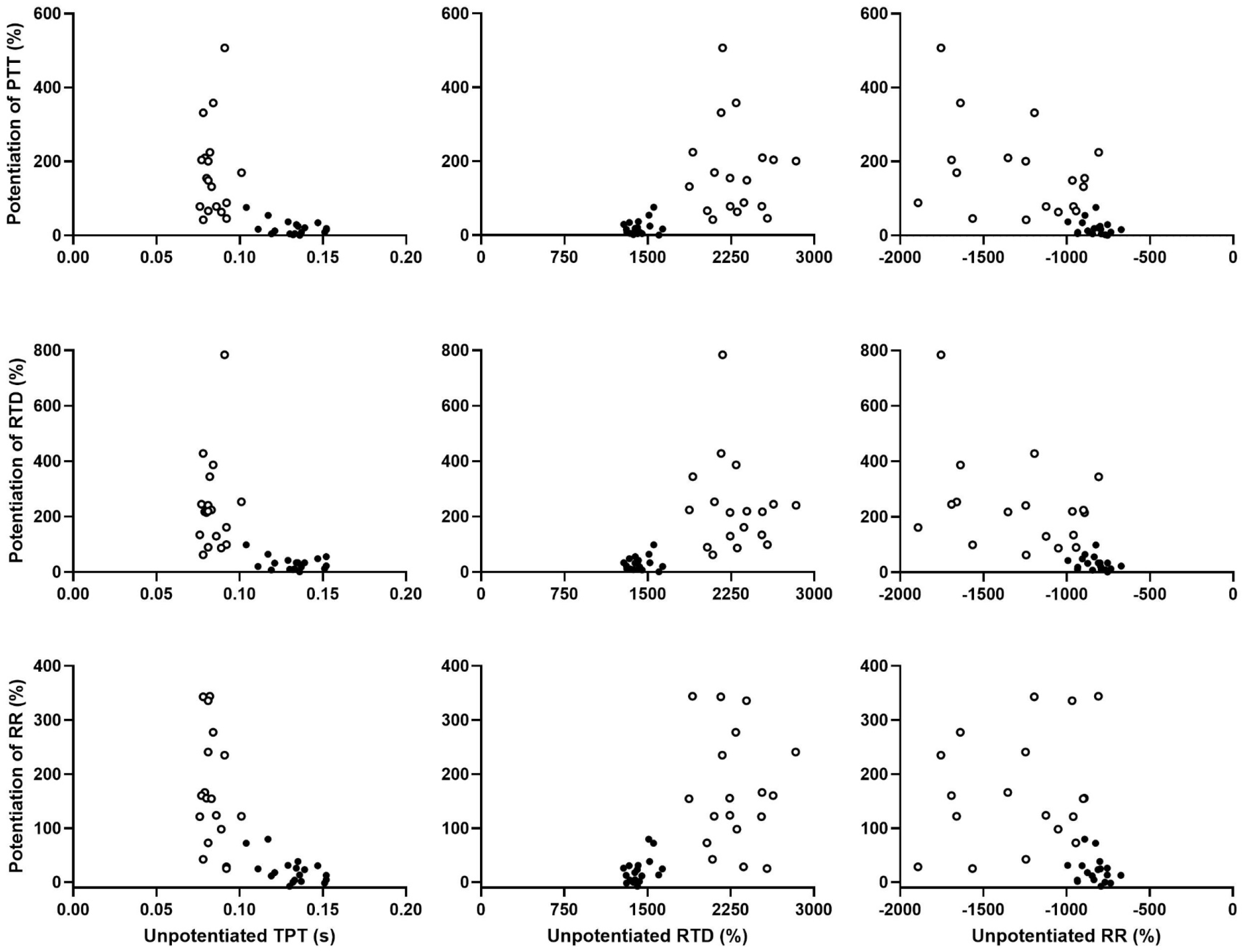
Relationship between relative changes in peak twitch torque (PTT, *top row*), rate of torque development (RTD, *middle row*), and rate of relaxation (RR, *bottom row*) and unpotentiated time to peak torque (TPT, *left column*), RTD, and RR in individual participants. Closed circles, triceps surae. Open circles, triceps brachii.

## Discussion

In numerous species, PAP occurs only in fast muscles, but whether a similar fiber type dependence exists in humans is still unclear. The purpose of the present study was to therefore to determine the effects of PAP on the twitch contractile properties of two muscles known to differ significantly in fiber type composition, i.e., the predominantly fast twitch triceps brachii versus the predominantly slow twitch triceps surae. We found that performance of a 6 s MVC increased PTT and enhanced the absolute and relative RTD in both muscle groups, but these effects were significantly larger in the triceps brachii. Prior contraction also shortened TPT and HRT and increased the absolute but not the relative RR in both muscle groups. These results support and expand current understanding of the fiber type-specific effects of PAP on the twitch contractile properties of human muscle.

At least two previous studies have observed greater increases in PTT following a brief MVC in faster versus slower muscles of humans (23,25). However, the effects of PAP on PTT have also been reported to be similar in the triceps brachii and the triceps surae (24), or even larger in the tibialis anterior (75% slow twitch; Ref. 32) versus the triceps surae (20). Although the reason for these discrepant findings is not clear, our results are comparable to the former studies (22,24), with the relative increase in PTT due to a 6 s MVC being >8-fold larger in the triceps brachii versus the triceps surae. Thus, in humans as in other species, PAP seems to increase twitch force to a much greater extent in fast muscle. Similarly, the greater increases in absolute and relative RTD that we observed in the triceps brachii versus the triceps surae are consistent with the results of animal studies indicating that PAP also increases the rate of force development primarily or exclusively in fast fibers (1). In contrast, previous studies of the effects of PAP in fast versus slow human muscles have either not measured twitch kinetics (23,25) or have only measured TPT and have found similar relative changes in both fast and slow muscles (20,24). PAP has a smaller effect on TPT than on RTD (29; Table 2), however, which may explain the latter findings.

In addition to the above-described changes in PTT, TPT, and RTD, we also found that PAP was accompanied by a shorter HRT and a faster absolute RR, with the latter effect being greater in the triceps brachii than in the triceps surae. In relative terms, however, there were no changes in RR in either muscle group. The latter results imply that PAP does not alter the fundamental determinant of relaxation, i.e., the rate of dissociation of myosin from actin (37), in either fast or slow human muscle, with the higher absolute RR simply reflecting the detachment of more such crossbridges per unit time. This conclusion is consistent with the fact that RLC phosphorylation does not alter the maximal rate of crossbridge cycling, as indicated by ATPase activity or unloaded shortening velocity (1). On the other hand, the shorter HRT in the potentiated state may be because twitch relaxation is a complicated process influenced by factors in addition to the rate of crossbridge detachment (37). For example, it has been suggested that PAP may accelerate relaxation via increases in inorganic phosphate during the conditioning contraction, thus favoring “backward” transition of still-attached crossbridges from their force-generating to their non-force-generating state (37). Alternatively, the greater PTT in the potentiated state may lead to more rapid “give” of the longest sarcomere in a myofibril and hence more rapid onset of the fast, i.e., “chaotic”, phase of relaxation (37,38).

The above results therefore support the conclusion that PAP influences the mechanisms of force development but not relaxation in human muscle in a fiber-type specific manner. Notably, however, no significant relationships were found between baseline twitch contractile properties and the extent of PAP on an individual basis, either within the triceps surae or triceps brachii or when data from both muscles were pooled together. These results contrast with previous studies that have reported modest-but-significant correlation between the increase in PTT due to PAP and, e.g., TPT in the unpotentiated state (25,26,30). Although the reason(s) for the lack of any such relationships in the present study cannot be determined, the absence of a strong correlation (including in previous studies (25,26,30)) indicates that factors other than fiber type (e.g., differences in tendon stiffness (19)) must also contribute to intraindividual differences in the effects of PAP.

As with all studies, there are limitations to the present investigation. The most obvious is the fact that muscle biopsies were not obtained to verify the fiber type distribution of the triceps surae and triceps brachii in the present participants. However, given the well-established differences in fiber type between these muscles (31,32) and the differing contractile properties (e.g., TPT) that were observed, this limitation seems minor. On the other hand, this may have contributed to the apparent lack of any relationship between individual muscle characteristics and the degree of PAP. Furthermore, despite clearly differing in fiber type both the triceps surae and triceps brachii are still mixed muscles, such that it is not possible to definitively ascribe our observations to strictly just slow twitch or fast twitch muscle fibers. We also did not quantify PAP in other muscles, at different joint angles/muscle lengths, in different states of fatigue, or during dynamic muscle contractions, all of which can influence the magnitude of the response (21,25,39-41). Likewise, we did not assess the effects of different conditioning contraction regimens, e.g., the duration and/or number of MVIC, which also affects the degree of PAP (20,29).

In summary, in the present study we tested the hypothesis that the effects of PAP on twitch contractile properties would be greater in the predominantly fast-twitch triceps brachii than in the predominantly slow-twitch triceps surae of humans. We found this to be true for PTT and absolute and relative RTD, but not for HRT or relative RR. We therefore conclude that, as in other species, PAP influences muscle contraction but not relaxation in humans in a fiber type dependent manner.

